# Evidence-based re-identification: Leveraging appearance cues to correct tracking mistakes rather than trying to prevent them

**DOI:** 10.64898/2026.11.08.765193

**Authors:** Vincent Coulombe, Quentin Leboulleux, Benoit Gosselin, Camille Testard, Benoit Labonté

**Affiliations:** CERVO Brain Research Centre, Université Laval, Québec, QC, Canada; Department of Psychiatry and Neuroscience, Faculty of Medicine, Université Laval, Québec, QC, Canada; Smart Biomedical Microsystems Laboratory, Quebec City, Canada; Department of electronics, Faculty of Science and Engineering, Université Laval, Québec, QC, Canada; Harvard Society of Fellows, Department of Molecular and Cellular Biology, Harvard, Cambridge, MA, U.S.A

## Abstract

Naturalistic behavioral neuroscience experiments, in which groups of animals interact in enriched environments over prolonged periods, require each animal’s identity to remain correct throughout the recording. Current multi-animal tracking tools assign identities frame by frame without revisiting past assignment decisions. A single, even sporadic, identity swap is therefore likely to persist until the end of the recording. Consequently, error-free tracking results become progressively less likely as recordings lengthen and cohorts grow. Even state-of-the-art multi-object tracking (MOT) methods that leverage appearance cues do so only to make each assignment more robust. This robustness, however, only delays the inevitable as even the most robust assignment process will eventually make a mistake over prolonged recordings. A mistake that will then persist just as it otherwise would have. Here, we introduce evidence-based re-identification, which uses appearance cues to correct tracking mistakes rather than prevent them. This fundamental shift allows the algorithm to build and progressively enrich appearance representations for each tracked subject. Tracking mistakes are then detected by continuously monitoring the evolution of these representations, triggering a correction whenever two representations drift toward one another in a sustained way. On two distinct, minutes-long benchmarks of mice interacting in enriched environments, motion-only trackers (DeepLabCut, SLEAP, ByteTrack) and the appearance-based BoT-SORT-ReID began accumulating identity errors within seconds, whereas ByteTrack paired with evidence-based re-identification remained reliable, outperforming even ByteTrack paired with the marker-based Tailtag system. This reliability was also positively correlated with the accuracy of the appearance extractor, but only when it was used through evidence-based re-identification. Foundation image encoders (CLIP, MegaDescriptor, DINOv3) showed promising out-of-the-box performance yet still required fine-tuning to be effective. Together, these results show that multi-animal tracking algorithms can remain reliable over prolonged periods only if they can reliably recover from tracking mistakes, and that fine-tuning appearance extractors remains essential to doing so.

## Introduction

Behavior not only reveals the strategies animals use to solve socio-environmental challenges, but also how these strategies fail in models of psychiatric disorders. Accurately quantifying behaviors is therefore an essential step towards understanding the neuronal mechanisms underlying cognition, emotion, and social interaction. Yet many of these strategies only emerge under naturalistic conditions (i.e., animals socially interacting within naturalistic environments over prolonged periods), which traditional protocols built around isolated animals or dyads in brief sessions are not designed to capture [1,2]. Recognizing these limitations, behavioral neuroscientists are progressively developing more naturalistic research protocols [3,4]. However, the very characteristics that make these protocols ethologically relevant also make them difficult to analyze [5,6]. Mapping ethological behaviors to their underlying neuronal mechanisms first requires knowing which animal is doing what, to whom, and when, which becomes increasingly difficult as groups grow larger and recordings longer. Because existing multi-animal tracking tools struggle to reliably maintain identities under these conditions [7], researchers fall back on manual observations, which are time-consuming and prone to observer bias [8, 9].

Over the past decade, deep learning-based tools such as SLEAP [7] and DeepLabCut [10,11] have made accurate, markerless multi-animal pose tracking accessible to behavioral neuroscientists. Both tools perform well over short recording sessions [7,11] but struggle to maintain correct identities when interacting animals are tracked over prolonged periods [6,7,12]. This limitation stems from how they link detections to identities over time as these associations are iteratively established between consecutive frames and are not revisited. As a result, assignment errors are likely to be carried forward rather than corrected. This means that a single, sporadic association mistake, such as two animals exchanging identities during a close interaction, is likely to persist for the remainder of the recording. With the probability of making such a mistake growing with both the frame count and the number of tracked subjects of a recording, error-free recordings quickly become the exception rather than the rule as experiments scale [7,12]. This leaves behavioral neuroscientists without useful tools to conduct ambitious ethological experiments in which biological conclusions hold only if each trajectory remains assigned to the correct animal throughout the entire recording. Consequently, for such experiments, the usefulness of a multi-animal tracking tool is not directly proportional to its reported accuracy on seconds-long recordings, but rather to its ability to produce accurate trajectories throughout entire recordings, a property we define as reliability.

Alternative tracking methods, such as marker-based approaches, are bypassing this limitation altogether by reading the animal’s identity directly from physical markers rather than iteratively updating their trajectories. This yields systems producing more reliable, but less spatio-temporally dense tracking results [3,4,13,14]. Notably, the Tailtag system [15] has shown promise for longitudinal studies by maintaining the identities of 20 mice over seven days in an enriched environment without performance degradation. Yet because this approach relies on markers suited to the anatomy of only a few rodent species, it cannot serve the full range of species studied in behavioral neuroscience.

The challenge of maintaining identities over time has also been studied extensively outside neuroscience, most notably in multi-object tracking (MOT). MOT methods aim to detect every relevant object in a scene and maintain a unique, persistent identity for each of them over time [16,17]. Most MOT methods follow the motion-based tracking-by-detection paradigm, in which identities are maintained by assigning each new detection to an existing trajectory, the same process used by DeepLabCut and SLEAP. Because modern object detectors are highly accurate [18–23], recent MOT contributions have largely focused on the assignment step, which is often framed as a data association problem [24,25]. However, standard MOT benchmarks consist of sequences usually lasting only a few seconds [17,26]. Therefore, the reliability over prolonged recordings of MOT methods have yet to be widely evaluated. Among motion-based MOT algorithms, ByteTrack [24] remains one of the strongest [25,27]. However, because it shares the same frame-by-frame assignment strategy as DeepLabCut and SLEAP, it is exposed to the same error-compounding problem, limiting its reliability over prolonged recordings.

A complementary line of MOT research is aimed at improving current motion-based methods by also leveraging the visual features of the tracked objects. This includes their colors, textures, shapes and even accessories such as clothing. These visual features, also referred to as appearance cues, are usually extracted from raw pixels using deep learning computer vision models called appearance extractors [28]. Appearance cues have been previously leveraged by MOT methods with great success, mostly to enhance the decisionmaking context of their detection to trajectory assignment process [29,30]. Concretely, appearance cues are enabling modern trackers to know both where the subjects are and what they look like. Consequently, most top-performing MOT algorithms are leveraging appearance cues in their assignment step [27,29,30]. Among them, BoT-SORT-ReID [27] stands out for its robustness in crowded scenes where its joint use of motion and appearance cues during association is shown effective at reducing identity switches between overlapping objects. Nevertheless, BoT-SORT-ReID only uses appearance cues to improve the robustness of assignment step itself, not to fundamentally change its tracking approach. Consequently, it shares the same practical limitation as DeepLabCut, SLEAP and ByteTrack.

Furthermore, trackers relying on appearance cues, such as BoT-SORT-ReID, can only be as reliable as the appearance cues themselves. Fortunately, the re-identification field has made substantial progress towards more capable appearance extractors [28]. A progress recently driven by computer vision foundational models, meaning large image encoders pretrained on vast and diverse datasets [31]. Many of these models are reported to produce meaningful appearance features without any fine-tuning [32–34]. First, BVRA’s MegaDescriptor [34] was introduced in 2023 as the first foundational model dedicated to animal re-identification, delivering state-of-the-art appearance extraction capabilities across a wide range of species. Second, OpenAI’s CLIP [33], trained on roughly 400 million image-text pairs, learned visual representations rich enough to transfer to new tasks, including re-identification, without fine-tuning [34]. Third, Meta’s DINOv3 [32], an image encoder pretrained in a self-supervised fashion on roughly 1.7 billion images, achieved state-of-the-art status across a wide range of visual benchmarks while keeping its backbone frozen, meaning no fine-tuning. Nevertheless, the re-identification capability of these appearance extractors has yet to be evaluated in behavioral neuroscience settings.

Here, we introduce evidence-based re-identification, a novel re-identification approach leveraging appearance cues to correct tracking errors rather than trying to prevent them. Moreover, it operates alongside any existing tracking algorithm without modifying it. When paired with the ByteTrack tracking algorithm, evidence-based re-identification is shown significantly more reliable than BoT-SORT-ReID, DeepLabCut, SLEAP, ByteTrack itself and even ByteTrack paired with the Tailtag system. We also evaluated the appearance extraction capabilities of the MegaDescriptor, CLIP and DINOv3 foundational image encoders in behavioral neuroscience settings.

## Results

We present evidence-based re-identification, a novel re-identification approach enabling multi-animal trackers to remain reliable over long recordings by monitoring how each subject’s appearance evolves over time to detect sustained identity swaps, a signal for tracking errors, and correcting them. Furthermore, the evidence-based re-identification method is completely self-contained and operates without modifying the tracking algorithm it is paired with (Figure 1a).

**Figure 1:**
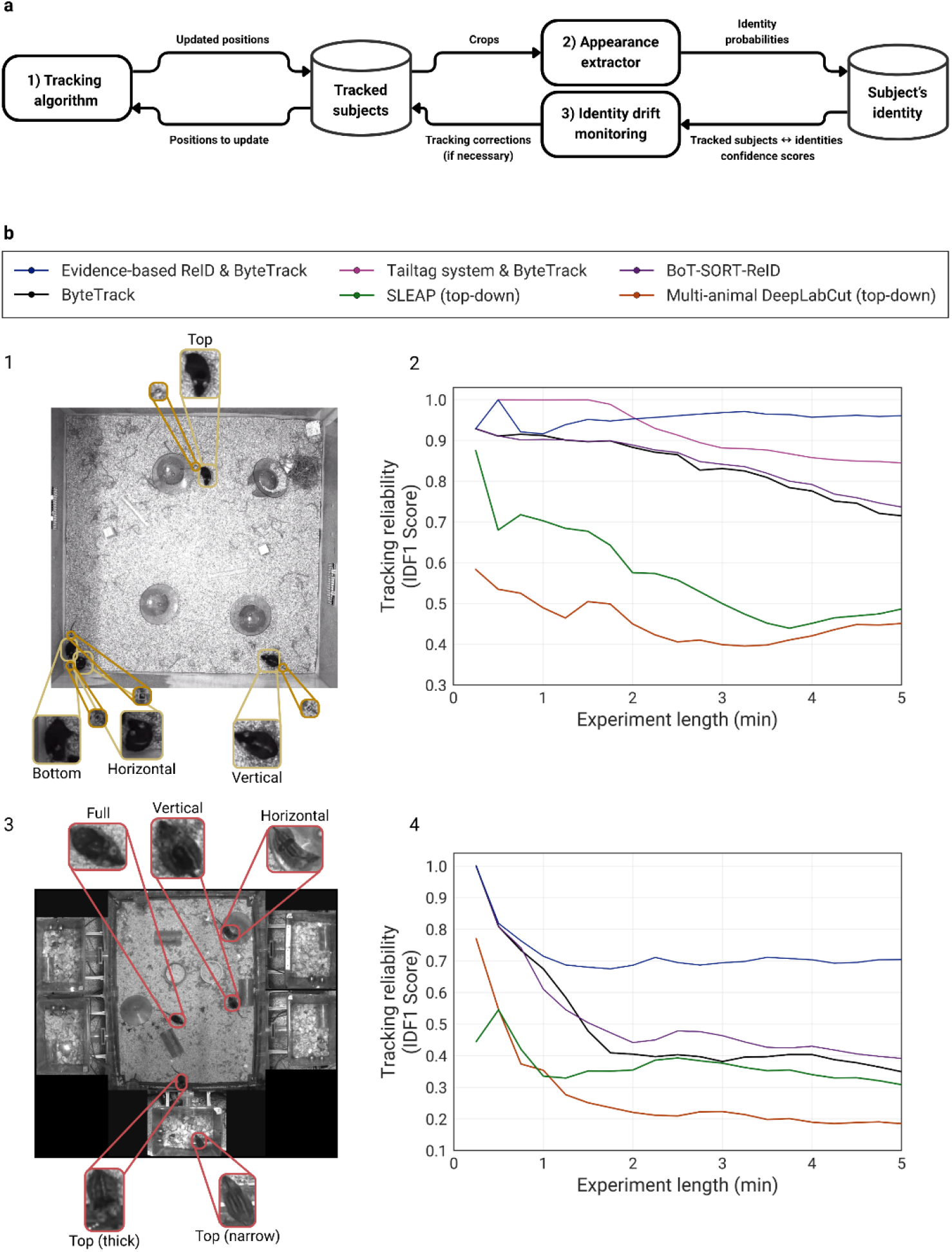
Method overview and tracking reliability results on two rodent benchmarks. **a**, Evidence-based re-identification overview. For every frame in a recording, the tracker follows each subject’s position while an appearance extractor assesses each subject identity (who each subject looks like). Rather than trusting the identity predictions of any single frame, the system accumulates this evidence over time to build a confident, established identity for each tracked subject. By monitoring these identities over time, the system can detect tracking errors are correct them. For instance, when two subjects start to progressively match each other’s established identity, their identity labels are swapped to correct the error. **b**, The evidence-based re-identification method was evaluated on two benchmarks of rodents freely interacting in open arenas. **1**, The MICE & ArUco tracking benchmark consists of 5-min recordings of four mice, each marked with a distinct shaved dorsal pattern (Top, Bottom, Horizontal, Vertical) and with a Tailtag marker for comparison. **2**, The evolution of the tracking reliability (the IDF1 score) over the course of the recording for each of method. Motion-based methods include ByteTrack (black), SLEAP (green) and DeepLabCut (orange). Re-identification-based methods include ByteTrack paired with evidence-based re-identification (blue), ByteTrack paired with Tailtag (pink) and BoT-SORT-ReID (purple). **3**, The StripedMice tracking benchmark consists of 5-min recordings of five mice, each marked with a distinct shaved dorsal pattern (Full, Top (thick), Top (narrow), Horizontal, Vertical). **4**, Same as 2, for the StripedMice benchmark. Tailtag is not included because the mice ate the markers.

Evidence-based re-identification is a recursive re-identification approach. Each frame, it gathers more identity evidence, progressively enriching the stored identity representations of each of the tracked subject. As shown in Figure 1a, this process starts whenever the paired tracker updates the subject’s positions. Then, an appearance extractor, estimates the probability of each subject to belong to each known identity. Finally, these estimations are accumulated over time, establishing an evolving identity representation for each subject. Consequently, by monitoring the evolution of the stored identity representations, the proposed method can detect tracking errors. More specifically, a tracking error is detected whenever the identity representations of two subjects drifted towards each others known identity beyond a configurable threshold. Then, a correction is triggered, and the identity labels of the two subjects are swapped back.

To evaluate the method under challenging conditions, we assembled two tracking benchmarks of rodents freely interacting in enriched open arenas containing bedding, running wheels, feeding trays, tubes, wooden blocks and other objects that frequently occlude the animals (Figure 1b). The first benchmark, MICE & ArUco, comprises four C57BL mice, each carrying a distinct shaved dorsal pattern (Top, Bottom, Horizontal, Vertical) as well as an ArUco marker from the Tailtag system [15]. The second benchmark, StripedMice, comprises five striped mice (*Rhabdomys pumilio*), each carrying a distinct shaved dorsal pattern (Full, Top (thick), Top (narrow), Horizontal, Vertical). In both benchmarks, animals are frequently occluded. Furthermore, each benchmark consists of 5-min (~8500 frames) recordings, a duration shown sufficient to determine if a multi-subject tracker remains reliable by measuring the evolution of the IDF1 tracking metric over time [12,15]. More specifically, a steadily declining IDF1 curve indicates that a tracker is making progressively more tracking error, meaning that they are compounding. In practice, this decline will tend to slow down and then stabilize around a particular score as the tracker stop making more mistakes. We refer to this score as a tracker steady state. As such, the higher tracker steady state is for a particular task, the more reliable it is for that task.

We compared the tracking reliability of six multi-animal tracking algorithms. Three motion-based tracking algorithms, meaning algorithms tracking subjects by leveraging only their positions and movements: ByteTrack [24] (black), top-down SLEAP [7] (green) and top-down multi-animal DeepLabCut [11] (orange). And three appearance-based tracking algorithms, meaning algorithms also leveraging appearance cues when tracking subjects: ByteTrack paired with evidence-based re-identification (blue), ByteTrack paired with the Tailtag system (pink) and BoT-SORT-ReID [27] (purple). As previously stated, their tracking reliability is quantified by the evolution of the IDF1 metric over the duration of the recordings (see *methods* for more details).

Evidence-based re-identification achieved the highest final IDF1 Score on both benchmarks (95% on MICE & ArUco and 71% on StripedMice; Figure 1b). As expected, every algorithm made tracking errors, but those unable to recover from them where either not even reaching steady state, meaning that their tracking errors were still compounding, or reached meaningless steady states. For example, DeepLabCut’s IDF1 score steadily fell below 20% (the IDF1 score obtained when assigning identities at random to 5 subjects) on the StripedMice benchmark. Even the appearance-based tracking algorithm BoT-SORT-ReID, which uses appearance to prevent tracking errors rather than correct them, could not reach a meaningful steady state in any of the benchmark. In fact, it performed only marginally more reliable than the motion-only tracking algorithm ByteTrack (73% versus 71% on MICE & ArUco and 39% versus 38% on StripedMice). In contrast, the two tracking algorithms able to detect and correct tracking errors (evidence-based re-identification and the Tailtag system) were both able to reach meaningful steady states. These results indicate that, over long recordings, using appearance to prevent tracking errors is not sufficient to achieve reliable tracking, but being able to detect and correct them is.

Furthermore, all tracking algorithms scored lower on the StripedMice benchmark than on the MICE & ArUco benchmark. This is due to the fact that the striped mice themselves contrast less with the arena background than the C57BL mice, making partially occluded animals harder to detect. The StripedMice benchmark recordings were also recorded at approximately 10 frames per second (rather than 25 for the MICE & ArUco benchmark). Consequently, the subjects in the StripedMice benchmark move 2.5 times farther between consecutive frames, often producing abrupt apparent jumps in position, which we term teleportations. Although undesirable in practice, these subpar tracking conditions make StripedMice a rigorous test as tracking algorithms able to reach a meaningful steady state on it are probably more likely to also hold up when used in other difficult real-world settings.

As previously mentioned, evidence-based re-identification relies on an appearance extractor to estimate who each subject looks like (Figure 1a). Here, an appearance extractor is defined as a pretrained deep learning computer vision model able to extract meaningful appearance cues from cropped images of each subject. These appearance cues can then be leveraged to predict a subject identity using two methods (see *methods* for more details). The first method is called open-set re-identification. Here, a subject’s identity is predicted by comparing its extracted appearance cues with a reference gallery containing multiple appearance cues from all the known subjects. The second method is called closed-set re-identification. Here, a linear classification head is trained on top of the extractor, enabling it to directly output a probability distribution over a fixed, predefined set of identities. Either way, we hypothesise that the practical reliability of the evidence-based re-identification method correlates with its appearance extractor’s re-identification capability, meaning its ability to accurately predict a subject identity.

Consequently, choosing the right appearance extractor is worthwhile endeavor. Currently, foundational computer vision models, pretrained on hundreds of millions of images, are increasingly used as general-purpose feature extractors as they usually require no task-specific fine-tuning [31-33]. Consequently, we evaluated the re-identification capabilities of the current state-of-the-art foundational computer vision models. More specifically, we selected BVRA’s MegaDescriptor [34] (trained specifically for animal re-identification), OpenAI’s CLIP [32] (trained on image–text pairs), and DINOv3 [33] (the current state of the art in self-supervised learning) in both its ViT and ConvNeXt variants (Figure 2a) for our experiment. Challenging the no task-specific fine-tuning claim, we evaluated them both frozen, meaning that their pretrained weights were unchanged, and after fine-tuning on the StripedMice-ReID dataset. This dataset consists of images from striped mouse recordings distinct from those in the StripedMice tracking benchmark.

**Figure 2:**
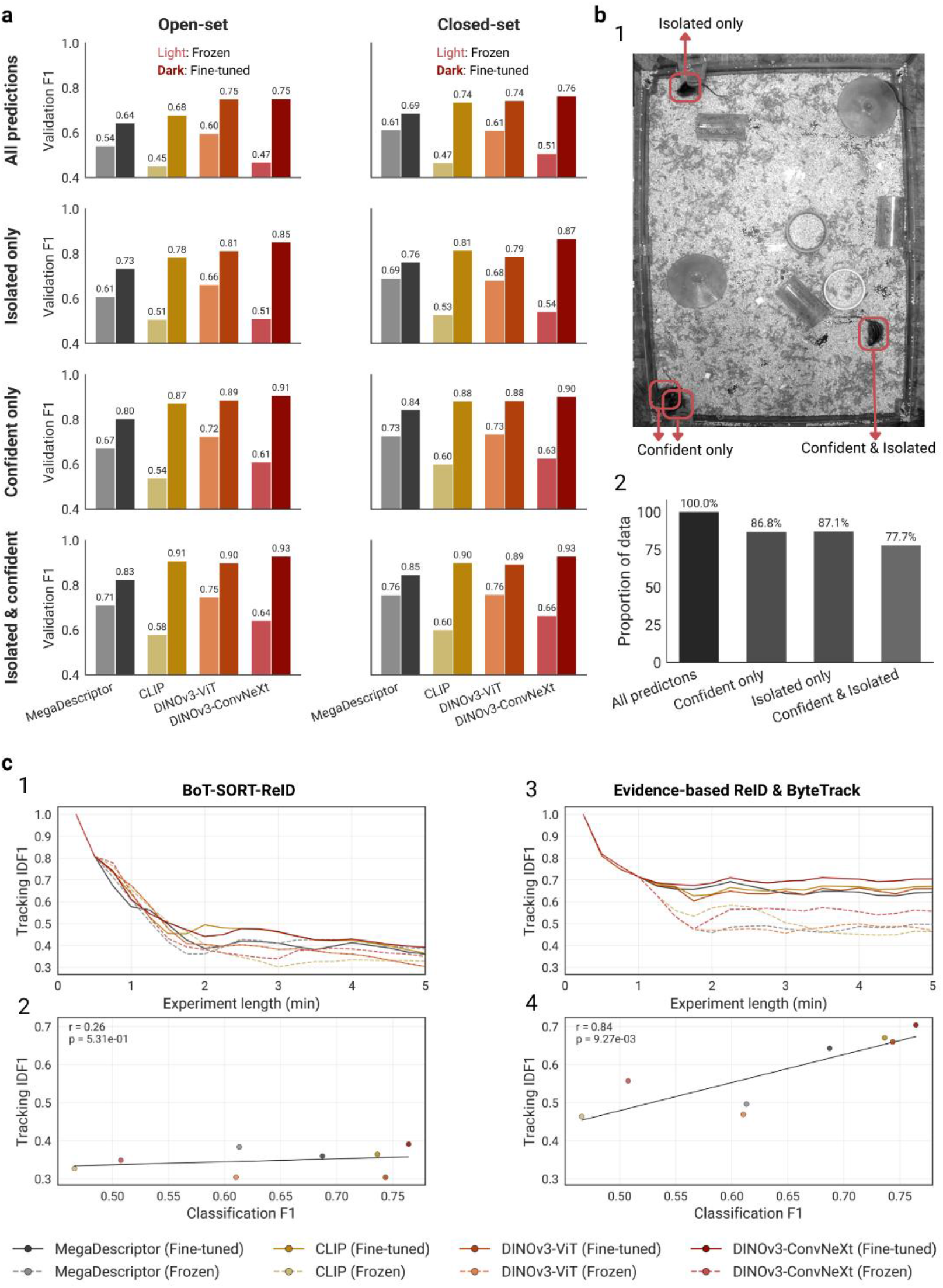
The impact of the appearance extractor re-identification capability and the re-identification method on downstream tracking reliability. Four foundational extractors are evaluated: BVRA’s MegaDescriptor [34] (grey), OpenAI’s CLIP [32] (gold), and Meta’s DINOv3 in its ViT (orange) and ConvNeXt (red) variants [33]. Fine-tuned extractors (dark) are trained on the training split of the StripedMice-ReID dataset, whereas frozen extractors (pale) retain their pretrained weights. Encoder capability is measured by F1 score on the StripedMice-ReID validation split, and tracking reliability by IDF1 on the StripedMice tracking benchmark. **a**, Identity prediction scenarios used to evaluate extractor capability. **1**, Illustration of the four scenarios: all visible subjects (all four bounding boxes), isolated subjects (top-left box), confident detections (the three boxes in the lower half) and isolated, confident detections (bottom-right box). **2**, Proportion of each scenario in the StripedMice tracking benchmark. **b**, F1 score of both the fine-tuned (dark) and the frozen (pale) variants of each extractor for each each re-identification scenario, using open-set classification (left) and closed-set classification (right). **c**, Effect of the re-identification method on downstream tracking reliability. **1**, IDF1 evolution of BoT-SORT-ReID when extracting appearances using each extractor from b. here, the appearances are extracted following the closed-set method. **2**, Correlation between the appearance extractor’s F1 score and the final IDF1 score achieved by BoT-SORT-ReID when leveraging it. **3, 4**, same as 1 and 2, for ByteTrack with evidence-based re-identification

Furthermore, a fair evaluation should also account for its task-specific decision-making context. Here, identifying a subject based on appearance cues alone can be more difficult in some scenarios as those cues can potentially be partially, or even fully, hidden. Such challenging conditions are common in the StripedMice benchmark, where the striped mice dorsal patterns are often hard to distinguish as they are occluded by either arena objects, other subjects or the subject’s own posture. To account for these challenging conditions in our experiment, we characterized the difficulty of each identity prediction using two proxy signals provided by the tracker itself. The first is the subject’s detection confidence score, which tend to reflect how visible the subject’s distinctive visual features are. For instance, subjects detected with lower confidence scores tend to be partially occluded or in an unusual posture. The second is isolation, as subjects in close proximity can occlude distinctive visual cues from one another. Taking into consideration these two signals yield four distinct appearance extraction decision-making scenarios. First, predicting the subject’s identities on all detections (all four bounding boxes in Figure 2a). Second, only when a subject’s detection is confident (all three boxes in the lower half). Third, when a subject is isolated only (top-left box). Fourth, only when a subject is isolated and its detection is confident (bottom-right box). Finally, the subjects in the StripedMice benchmark fall in the fourth scenario category during more than 75% of the recordings (Figure 2b).

The experiment itself consisted of evaluating the re-identification capability of the selected appearance extractors by measuring their identity classification F1 scores on the StripedMice-ReID validation split. Each extractor re-identification capability was evaluated on all four, previously described, decision-making scenarios as well as when using both the open-set and the closed-set classification method (Figure 2a). Overall, no significant gap between the open-set and the closed-set methods were observed. More notably, fine-tuning proved essential. The best extractor, fine-tuned DINOv3-ConvNeXt, achieved a closed-set F1 score of 76% on all detections and 93% on isolated and confident detections, whereas the best frozen extractor (DINOv3-ViT) reached only 61% and 76%, respectively. We attribute this gap to two factors. First, neuroscience recordings differ markedly from the internet images these foundational extractors were pretrained on. Here, subjects are relatively small, often occluded, filmed from above and the frames are in grayscale. Second, these extractors are mostly trained to distinguish between object categories, a much more trivial task than distinguishing individuals of the same species differing only subtly in appearance. consequently, even state-of-the-art foundational vision encoders must be fine-tuned on the target data distribution to reliably re-identify subjects in neuroscience experiments.

We also evaluated the impact of the re-identification capability of the appearance extractors on the downstream reliability of two tracking algorithms (Figure 2c). More specifically, we measured the tracking reliability of the BoT-SORT-ReID (right) and the ByteTrack paired with evidence-based re-id tracking (left) algorithms by measuring the evolution of their achieved IDF1 scores over time on the StripedMice benchmark. Notably, no matter the extractor used, BoT-SORT-ReID failed to reach a meaningful steady state, with a best final IDF1 of 39% when paired with the DINOv3-ConvNeXt extractor (the result of this particular pairing is also reported in Figure 1b). Furthermore, no significant correlation (r=0.26; p=5.31e-1) between the re-identification capability of the appearance extractors and the tracking reliability of the BoT-SORT-ReID tracking algorithm leveraging them were found, indicating that increasing an extractor’s identity-predicting accuracy is not guaranteed to increase the reliability of the tracking algorithm leveraging it. This lack of correlation also emphasizes that the inability of the BoT-SORT-ReID algorithm to reach a meaningful steady state in both benchmarks is likely not due to a shortcoming in its appearance extractor’s capability, but instead due to the way the extracted cues are leveraged. On the other hand, evidence-based re-identification was able to reach steady state no matter the appearance extractor it used, achieving a final IDF1 of 71% when paired with the DINOv3-ConvNeXt extractor (the result of this pairing is also reported in Figure 1b). Furthermore, a positive correlation (r=0.84; p=9.27e-3) between the leveraged appearance extractor re-identification capabilities and the downstream tracking reliability is observed, confirming our initial hypothesis. Notably, no frozen appearance extractors were able to lead to a tracking algorithm reaching an IDF1 score above 60% while the tracking algorithms powered by fine-tuned appearance extractors all reached a final IDF1 score of at least 60%, further supporting the claim that even the current state-of-the-art foundational extractors need to be fine-tuned to produce reliable trackers.

Therefore, we are assessing that even the current state-of-the-art image encoders need to be fine-tuned to power reliable behavioral neuroscience multi-animal tracking algorithms. Consequently, researchers aiming at leveraging such tools will, unfortunately, have to gather a fine-tuning dataset specific to their needs. As such, they will have to collect, curate and label subject identity data points. A task easier said than done as an effective machine learning algorithm need to be fine-tuned on datasets containing diverse and challenging set of examples, as well as being representatives of the predicting conditions [35]. Concretely, in the behavioral neuroscience context, this means gathering data points of either the tracked subjects themselves or subjects with the same distinguishable visual features (i.e. dorsal shaving patterns). This also means that these data points must include examples of all these subjects in as many poses and as many occlusion level as possible. Here, we pose the hypothesis that the most effective way to achieve such a dataset in practice is not to simply increase the amount of collected data at once, as this could mean gathering a lot of redundant data points (i.e. a subject remaining immobile during the whole data collection process). Instead, we hypothesize that for a data collection process to yield a diverse dataset, it must contain data points taken from many segmented recording sessions.

Figure 3a show that there is a positive correlation between the tracking reliability of the ByteTrack & evidence-based re-identification tracking algorithm and the capability of the appearance extractor powering it. This holds for both of our benchmarks (r=0.61; p=7.15e-05 and r=0.34; p=4.45e-02 on the MICE & ArUco and StripedMice benchmarks respectively). Figure 3b and Figure 3c show that this correlation grows stronger as the amount of recording sessions in the fine-tuning dataset increases. More specifically, the correlation coefficient goes from (0.54; p=1.68e-2) to (0.78; p=2.13e-04) on the MICE & ArUco benchmark and from (0.28; p=1.64e-01) to (0.67; p=4.97e-02) on the StripedMice benchmark when the amount of recording sessions in the fine-tuning dataset goes from less than four to more than four. Furthermore, Figure 3d shows that appearance extractors fine-tuned on highly ablated versions of both fine-tuning datasets, down to about 13% and 17% of their original volume for the MICE and StripedMICE identity datasets respectively, could still lead to reliable tracking systems. In order words, randomly removing more than 80% of the data points in the identity dataset on which an appearance extractor was fine-tuned on did not impact the reliability of the tracking algorithm leveraging it. This indicates that identity datasets tend to be filled with redundant noise. Although, we note an increase in the variability of the measured downstream tracking performance as the dataset volume decreases. This suggests that, as datasets got smaller, its noise over signal ration diminished, increasing probability of randomly removing useful data points. Furthermore, Figure 3e shows that appearance extractors fine-tuned on datasets containing four or more recording sessions powered significantly more reliable tracking algorithms than those fine-tuned on datasets containing four or fewer recording session. Overall, Figure 3d and Figure 3e indicate that appearance extractors fine-tuned on bigger datasets are not guaranteed to power more reliable tracking system, but those fine-tuned on datasets containing significantly more recording session are significantly more likely to, supporting our initial hypothesis.

**Figure 3:**
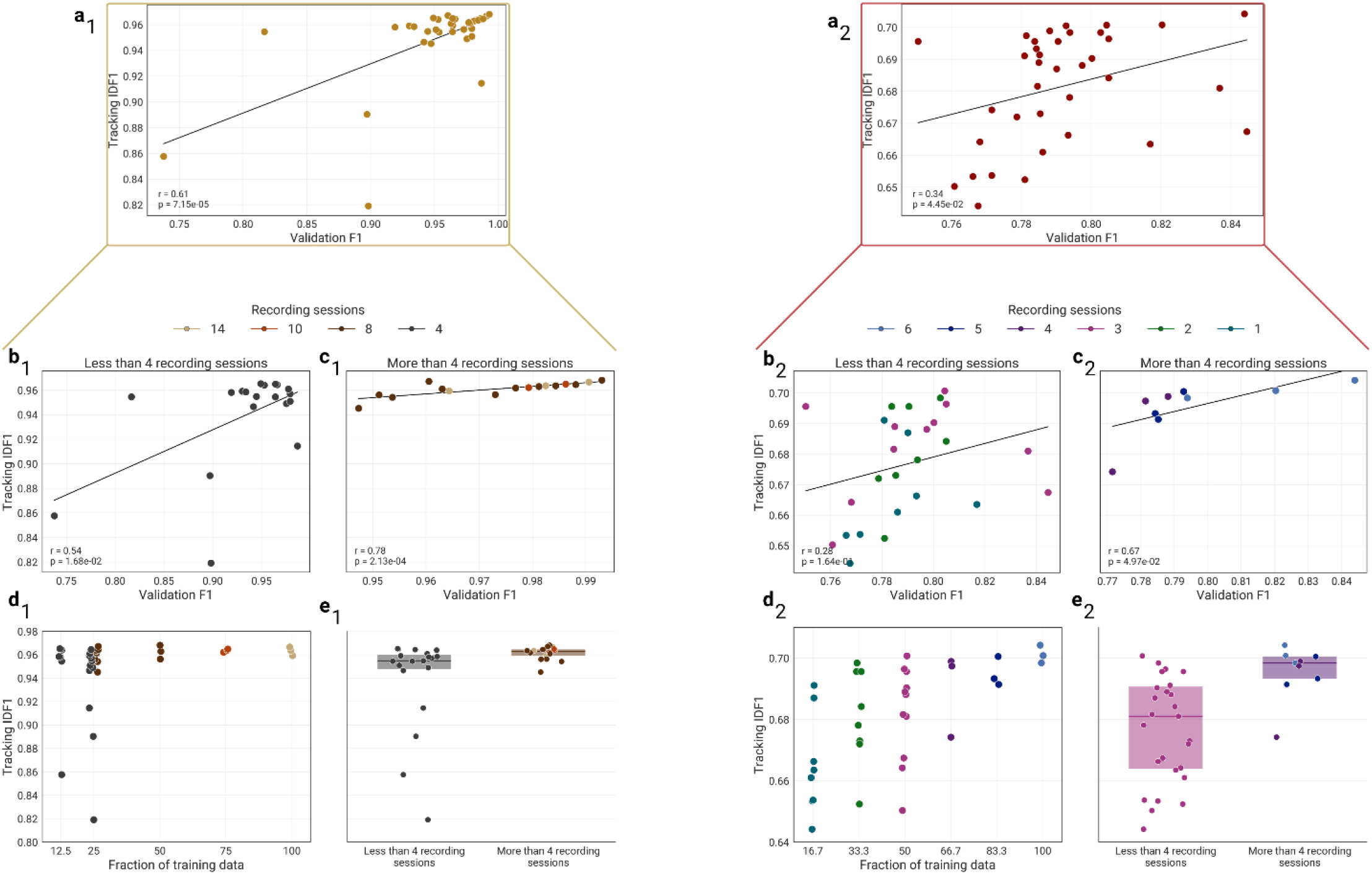
Impact of fine-tuning data quantity and diversity on the fine-tuned appearance extractor re-identification generalizability and downstream tracking reliability. In all panels, ByteTrack is paired with the evidence-based re-identification method leveraging DINOv3-ConvNeXt appearance extractors fine-tuned on datasets of varying size and diversity. Encoder generalizability is measured by the F1 score and tracking reliability by the IDF1 score. Each point represents one fine-tuned extractor. All subpanel 1 show F1 scores on the MICE shaving-patterns dataset and IDF1 scores on the MICE & ArUco tracking benchmark while all subpanel 2 show the same metric scores for extractors fine-tuned on the StripedMice-ReId dataset and evaluated on the StripedMice tracking benchmark. **a**, Correlation between the extractor F1 score (x-axis) and the tracking IDF1 score (y-axis) across all extractors. **b**, subset of a containing only the extractors fine-tuned on four or fewer recording sessions. **c**, subset of a containing only the extractors fine-tuned on more than four recording sessions. **d**, IDF1 score as extractors are fine-tuned on progressively more data (x-axis) and recording sessions (color). **e**, Effect of fine-tuning data diversity on downstream tracking reliability: IDF1 score for extractors fine-tuned on four or fewer (left) versus more than four (right) recording sessions.

## Discussion

Currently, the most best performing Multi-Object-Tracking (MOT) algorithms, including BoT-SORT-ReID, follow the same re-identification approach of enhancing motion-only tracking algorithms by also leveraging appearance cues when assigning newly detected subjects to existing trajectories [27,30] through a process known as object re-identification [28]. In other words, the current dominant approach in MOT is to leverage extracted visual features to minimize the probability of making tracking errors. This strategy is dominant as it is well suited to current MOT benchmarks, which consist of short, densely populated video clips (about 25 seconds on average [17,26]) and therefore directly reward methods that make the fewest assignment errors. As a result, new MOT methods are constrained to follow the currently established re-identification approach to remain competitive on these benchmarks.

However, many real MOT applications have requirements completely different from what these benchmarks are rewarding. For instance, in behavioral neuroscience, researchers need multi-animal tracking tools generating the data on which their conclusions rest to maintain the integrity of each subject’s trajectory. In other words, they are far more concerned about a multi-animal tracking tool’s ability to remain reliable over prolonged periods than its accuracy on short clips.

As such, to properly evaluate the reliability of modern MOT and multi-animal tracking methods, we assembled two behavioral neuroscience MOT benchmarks specifically designed to evaluate the reliability of tracking algorithms. Both benchmarks feature freely interacting rodents in enriched open arenas containing bedding, running wheels, feeding trays, tubes, wooden blocks and other objects that frequently occlude the animals, and their videos last minutes rather than seconds. Our experimental results show that this temporal discrepancy is crucial in discriminating between algorithms remaining reliable over prolonged periods from those that do not. More specifically, by monitoring the evolution of the IDF1 tracking metric over time, we were able to assess the rate at which each of the evaluated tracking algorithms makes permanent tracking errors. As such, a continuously decreasing IDF1 score indicates that a tracking algorithm is making progressively more mistakes, which happens whenever mistakes are made at a higher rate than they are recovered. The tracker then enters a degenerate state in which errors compound on top of each other, making it progressively less reliable. For instance, all the benchmarked motion-only tracking algorithms (DeepLabCut [11], SLEAP [7] and ByteTrack [24]) systematically entered this degenerate state within a few seconds of tracking. This IDF1 decline stops only once them metric calculation and the mistake-to-recovery ratio stabilizes. Then the tracker reached steady state. In some cases, such as DeepLabCut on the StripedMice benchmark, this steady state is so low that randomly assigning identities at every frame would have provided better reliability. Overall, these results indicate that relying on motion alone is not sufficient to guarantee reliable multi-animal tracking over prolonged periods.

Furthermore, our results show that the currently dominant MOT strategy of enriching the detection to trajectory association step with visual cues does not guarantee reliable multi-animal tracking over prolonged periods. For instance, the visual cues and motion-based BoT-SORT-ReID [27] algorithm performed only marginally better than the motion-only ByteTrack algorithm on both benchmarks, indicating that the re-identification enhancement did little to improve reliability. In other words, the extracted appearance cues did not prevent the tracker from making mistakes. We therefore hypothesized that the only way for a tracking algorithm to remain reliable over prolonged periods is to reliably correct the mistakes it will inevitably make. We developed the evidence-based re-identification approach to test this hypothesis. The proposed idea is to switch from a mistake-mitigation mindset to a mistake-correction mindset, meaning focusing on correcting tracking errors rather than preventing them. This shift allowed us to address three major limitations preventing tracking algorithms leveraging the current established MOT re-identification approach from remaining reliable over prolonged periods.

First, leveraging appearance cues within the detection to trajectory assignment process itself forces the algorithms to extract appearance cues from detections that have not yet been assigned, some of which may be false positives. Consequently, this prohibits the algorithm from refining its appearance extraction decision-making context as the spatial location of the subjects have yet to be accurately assessed. While this is not a meaningful limitation in most use cases, it becomes problematic when considering that the appearance cues can only meaningfully enhance an assignment decision when accurate. Furthermore, our results show that this the extracted appearance cues can become almost 20% less accurate when multiple subjects are closely interacting. algorithms following the established approach are therefore inevitably relying on appearance cues of varying accuracy. In contrast, evidence-based re-identification takes place after the detection to trajectory assignment process. As such, its inputs are the coordinates of tracked subjects rather than those of detected entities. This allows it to account for the spatial status of every subject and considers only appearance cues extracted in trustworthy decision-making scenarios.

Second, relying on appearance cues extracted from detections rather than trajectories removes the temporal dimension from the assignment decision-making context. This is because an appearance cue cannot be placed in temporal context before it has been assigned to a trajectory. Therefore, the algorithms are forced to hypothetize that the cues extracted from detections are accurate. Yet our results show that even fine-tuned, state-of-the-art appearance extraction models inevitably make mistakes. Combined, these two facts create a significant limitation as any sporadic appearance extraction mistake can cause a detection to trajectory assignment error, starting the very error-compounding degenerate state the re-identification approach was trying to prevent in the first place. In contrast, evidence-based re-identification, as its name implies, make correction decisions based on progressively enriched stored identity representations. As a result, extracted appearance cues must drift in a sustained manner to trigger a tracking correction, filtering sporadic extraction mistakes by design.

Third, since both previous limitations make tracking errors inevitable, a tracking algorithm must include a robust mechanism to recover from them to even have a chance of remaining reliable over prolonged periods. Our results support this claim: the only two algorithms that reached a meaningful steady state were ByteTrack paired with evidence-based re-identification and ByteTrack paired with the Tailtag system, which are also the only two algorithms with a robust mistake-correction strategy. In contrast, BoT-SORT-ReID was only marginally more reliable than ByteTrack despite leveraging appearance cues, which shows that appearance information alone cannot compensate for the absence of a correction mechanism. Overall, our results confirm our initial hypothesis that tracking algorithms can remain reliable over prolonged periods only if they can reliably recover from tracking errors.

Our results also show a strong, positive correlation between the accuracy of an appearance extractor and the reliability of the tracking algorithm leveraging it, but only when the extractor is used through evidence-based re-identification. This indicates not only that evidence-based re-identification uses extracted appearance cues more effectively than other MOT re-identification approaches, but also that improving the capability of its appearance extractor is a worthwhile endeavor as it can translate into more reliable tracking.

Regarding extractor accuracy, we evaluated four appearance extractors: OpenAI’s CLIP, BVRA’s MegaDescriptor and both Meta’s DINOv3 (ConvNeXt and ViT). All are promoted as foundation, plug-and-play feature extraction models, which in our context implies that they should provide accurate appearance extraction out of the box. Yet our results show that all four require fine-tuning to reach a satisfactory F1 identity classification score. We attribute this to the unusual data distribution of our benchmarks. Top-down, black-and-white, low-saturation, zoomed-in crops of interacting mice bearing shaved dorsal patterns fall well outside the natural-image distribution these models were pretrained on. Given how unreliable deep-learning models can be on out-of-distribution data, it is notable that the frozen backbones performed as well as they did. As such, we interpret their frozen F1 scores as evidence of the generalization ability of these foundational models rather than against it.

Since fine-tuning is necessary, the question becomes how to gather a diverse and representative fine-tuning dataset as efficiently as possible. On that front, our dataset ablation experiment shows that not all fine-tuning datasets are equally effective. More specifically, we found that identity datasets created from longitudinal data are filled with redundant data points. In fact, we found that appearance extractors fine-tuned on as little as 12.5% of the available identity data powered tracking algorithms just as reliable as those fine-tuned on all of it. In contrast, our results show that appearance extractors fine-tuned on datasets built from more recording sessions powered substantially more reliable tracking algorithms. We interpret these results as indicating that the most efficient way to increase the diversity of an identity dataset is to sample from more recording sessions rather than to collect more data points within the same sessions. However, this conclusion becomes more nuanced on more challenging benchmarks. For instance, the correlation between the accuracy of the fine-tuned appearance extractor and the tracking reliability of the tracking system leveraging it was much weaker on the more difficult StripedMice tracking benchmark. This suggests that task difficulty may moderate how much a more accurate appearance extractor improves tracking reliability. Exploring this relationship seems to be a promising direction for future research.

Finally, the fact that the current state-of-the-art foundational image encoders still need fine-tuning in order to power reliable behavioral neuroscience multi-animal tracking algorithms points to a clear and potentially impactful future research direction: the creation of a behavioral neuroscience foundational dataset. Such a dataset could define a standard dictionary of shaved dorsal patterns and provide diverse and representatives data points for each, enabling the fine-tuning of a specific foundational encoder to achieve satisfactory appearance extraction capabilities out of the box, provided the standard is respected. To assemble it, we encourage behavioral neuroscience researchers to join forces and pool data spanning multiple rodent species and tracking environments.

## Methods

### Multi-animal tracking

At a high level, multi-animal tracking can be formally defined by letting 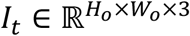 denoting the frame at time *t* and let the tracker emit a set of detections *T*_*t*_.

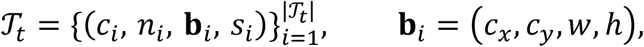

where *c*_*i*_ is the predicted specie, *n*_*i*_ *∈* ℤ the subject id, **b**_*i*_ the bounding box in centre form and *s*_*i*_ *∈* [0,1] the detection confidence. Each tracked subject is addressed by the unique key *u* = (*c, n*), and the *M* keys expected in the recording are enumerated once at initialisation, giving an index map *u* ↦ *j ∈* {1, …, *M*}.

### Determining re-identification priorities

Figure 2a showed that re-identification models are more reliable on isolated subjects. Furthermore, Figure 1 of the supplementary material showed that the re-identification (validation) process is a latency bottleneck for the whole tracking pipeline. Therefore, we choose to restrict the pipeline to re-identify the subset *S*_*t*_ ⊆ *T*_*t*_ of *isolated* subjects reported by the tracker.

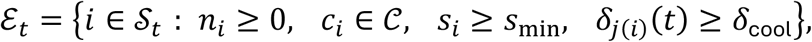

where *C* is the set of validated species, *s*_min_ the minimum detection confidence threshold, and *δ*_*j*_(*t*) *∈* ℕ counts the frames elapsed since track *j* was last examined. The cooldown *δ*_cool_ prevents redundant calculations, as high throughput trackers will produce sequential detections with very similar pixel content.

At most *B* candidates are processed per frame. Candidates are ranked by a priority based on both detection quality and staleness. More specifically, the *B* highest-priority elements are drawn from a max-heap, *ℬ*_*t*_ = top − *B*(*ℰ*_*t*_, *π*). Note that *δ*_*j*(*i*)_(*t*) is normalized by the staleness horizon *δ*_max_. Therefore, a confident but recently verified tracking bounding box will have a lower priority score than another that has not been observed for a long time. The per-frame latency of the re-identification pipeline is thus *O*(*B*) and independent of *M*, making it configurable.

### Re-identification model

For each selected candidate the bounding box is dilated by a factor *γ* > 1, 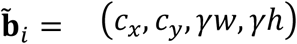, clipped to the image and cropped. Crops are resized to the network input resolution *h* × *w*, converted to RGB and normalised. The resulting batch 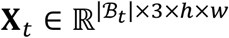 is passed through the re-identification backbone *ϕ* and its classification head *ψ*,

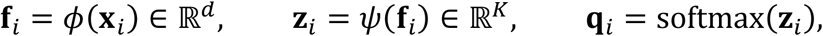

yielding a per-crop identity probability distribution **q**_*i*_.

### Rolling identity evidence

We maintain, for every tracked subject *j*, an identity evidence state **p**_*j*_(*t*) *∈* Δ^*K*−1^ recursively updated as an exponential moving average,

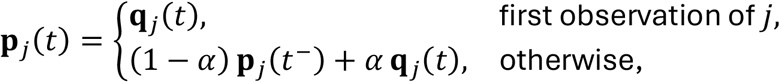

where *t*^−^ denotes the previous frame at which *j* was examined and *α ∈* (0,1) is the adaptation rate. Unrolling the recursion,

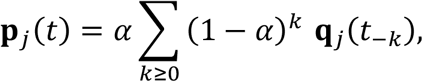

shows that the identity evidence is, concretely, a weighted average over the last ≈ 1/*α* observed identities of subject *j*, yielding dynamically updated subjects to identities confidence scores.

### Identity drift monitoring

Disabled identities are first excluded following:

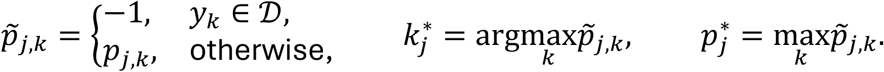

Furthermore, a prediction is *confirmed* when its updated confidence score exceeds

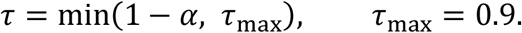

A single observation can raise a component of **p**_*j*_ by at most *α*, so requiring 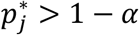 makes confirmation impossible from one frame alone and enforces temporal consensus, while *τ*_max_ caps the requirement for aggressive smoothing regimes (*α* → 0).

More than one tracked subject claiming the same identity within one frame signal an unresolved ambiguity rather than a correction. Let 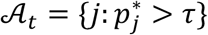 be every identity claimed more than once,

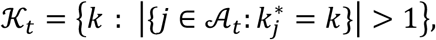

Gives the retained set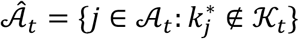.

Finally, identity confirmations are accumulated in a matrix **H** *∈* ℝ^*M*×*K*^. For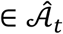,

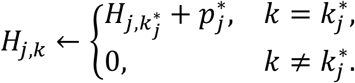

Zeroing the remainder of the row makes the evidence strictly consecutive, meaning that every identity confirmation resets the accumulated evidence for *j*. This mechanism ensures that successive corrections are separated by a full re-accumulation period.

### The IDF1 metric

The IDF1 metric [36] evaluates a tracking algorithm’s ability to maintain the correct identities of tracked objects over time by reporting the fraction of detections that carry the correct, temporally consistent identity over a sequence.

IDF1 rests on a one-to-one correspondence between ground-truth and predicted trajectories. Let *g* = {*τ*_1_, …, *τ*_*M*_} be the set of ground-truth identities and *P* = {*γ*_1_, …, *γ*_*N*_} the set of predicted identities. For a candidate pairing (*τ, γ*), we count the number of frames in which both are present and their bounding boxes coincide, i.e. their intersection-over-union exceeds a threshold *θ* (0.5),

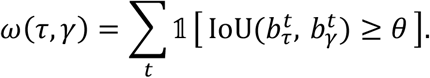

Each ground-truth identity may be linked to at most one predicted identity, and vice versa. The optimal assignment *X*^*⋆*^ *∈* {0,1}^*M*×*N*^ is the one that maximises the total number of correctly identified detections,

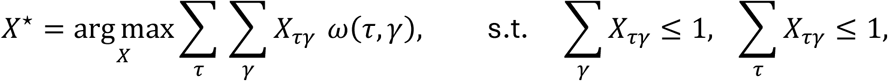

a bipartite matching solved with the Hungarian algorithm. Under this global assignment, three quantities are defined:

- **IDTP** (identity true positives): predicted detections assigned to the correct identity: 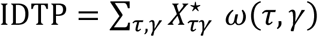;
- **IDFN** (identity false negatives): ground-truth detections that received a wrong identity or none: IDFN = |*D*_*g*_| − IDTP;
- **IDFP** (identity false positives): predicted detections that do not match their assigned ground-truth identity: IDFP = |*D*_*P*_| − IDTP,

where |*D*_*g*_| and |*D*_*P*_| are the total numbers of ground-truth and predicted detections in the sequence. Identification precision and recall follow directly,

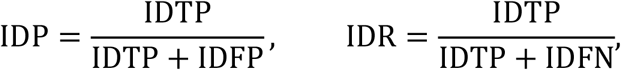

and IDF1 is their harmonic mean, equivalently written in terms of the three counts alone:

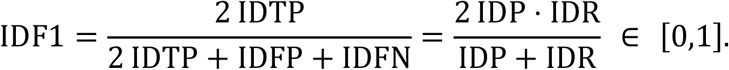

### Closed-set re-identification

Closed-set re-identification happens when the identity label space is fixed and identical between training and inference, meaning that only the identities present in the training dataset are expected to also be present in the test dataset:

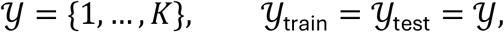

### Open-set re-identification

Open-set re-identification happens when identities present at inference are not constrained to those seen during training. The test-time gallery may contain novel subjects, and a query may correspond to no gallery identity at all,

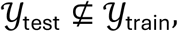

An opened-set re-identification model is often achieved by first training a model to cluster together appearances belonging to the same individuals, a type of training regimen commonly known as Metric Learning [37]. More specifically, Metric Learning algorithms learns to optimize a contrastive objective *ϕ* so that embeddings of the same identity are grouped while embeddings of different identities are far apart. Embeddings are ℓ_2_-normalised onto the unit hypersphere,

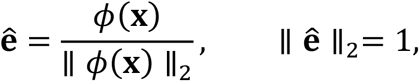

so that comparison reduces to the cosine similarity between two embeddings ê_*a*_, ê_*b*_,

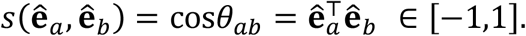

The established identity appearances are represented by an identity database commonly referred to as a gallery (*g*),

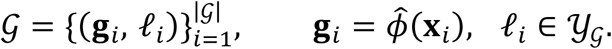

During inference, the system classifies a new identity **x**_*q*_ by first calculating its embedding ê_*q*_ then performing a *k* nearest neighbor classification using the cosine similarity as distance metric. The identity is decided by a majority vote over those neighbors, subject to a rejection threshold *γ* on the best match,

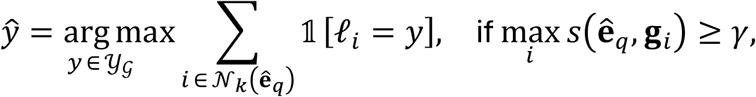

