## Supplemental Figure 1 for "Evidence-based re-identification: Leveraging appearance cues to correct tracking mistakes rather than trying to prevent them": Supplemental Figure 1.pdf

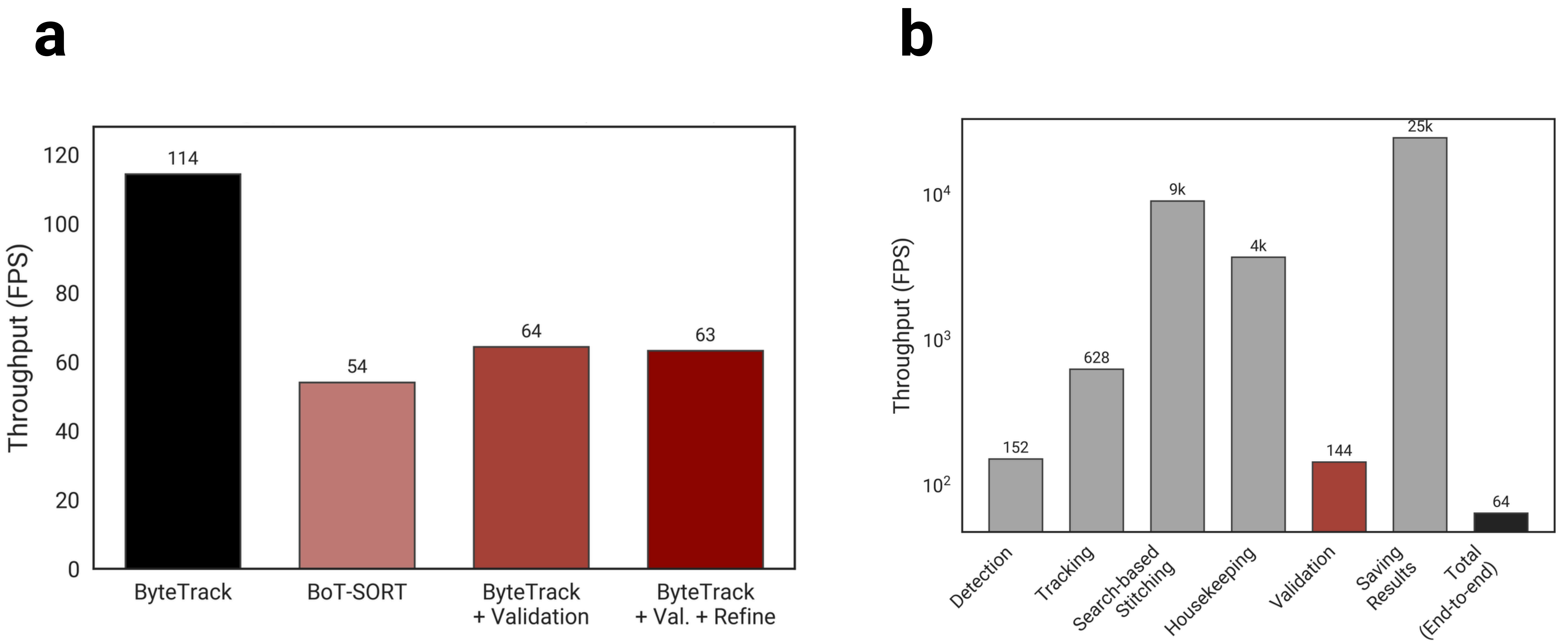

a
b
Throughput (FPS)
120
100
80
60
40
20
0
114
54
64
63
ByteTrack
BoT-SORT
ByteTrack
+ Validation
ByteTrack
+ Val. + Refine
Throughput (FPS)
$10^4$
$10^3$
$10^2$
152
628
9k
4k
144
25k
64
Detection
Tracking
Search-based
Stitching
Housekeeping
Validation
Saving
Results
Total
(End-to-end)
